# Genetic Adaptation of *C. elegans* to Environment Changes I: Multigenerational Analysis of the Transcriptome

**DOI:** 10.1101/194506

**Authors:** İrem Çelen, Jung H. Doh, Chandran R. Sabanayagam

## Abstract

Abrupt environment changes can elicit an array of genetic effects. However, many of these effects can be overlooked by functional genomic studies conducted in static laboratory conditions. We studied the transcriptomic responses of *Caenorhabditis elegans* under single generation exposures to drastically different culturing conditions. In our experimental scheme, P_0_ worms were maintained on terrestrial environments (agar plates), F_1_ in aquatic cultures, and F_2_ back to terrestrial environments. The laboratory N2 strain and the wild isolate AB1 strain were utilized to examine how the genotype contributes to the transcriptome dynamics. Significant variations were found in the gene expressions between the “domesticated” laboratory strain and the wild isolate in the different environments. The results showed that 20% - 27% of the transcriptional responses to the environment changes were transmitted to the subsequent generation. In aquatic conditions, the domesticated strain showed differential gene expression particularly for the genes functioning in the reproductive system and the cuticle development. In accordance with the transcriptomic responses, phenotypic abnormalities were detected in the germline and cuticle of the domesticated strain. Further studies showed that distinct groups of genes are exclusively expressed under specific environmental conditions, and many of these genes previously lacked supporting biochemical evidence.

## Introduction

All living systems possess the fundamental property to regulate physical or genetic states in response to environmental stimuli. *Caenorhabditis elegans* (*C. elegans*) offers one of the unique models to study transgenerational adaptation to environmental changes as it can reproduce in large numbers and has a natural ability to live in both terrestrial and aquatic conditions (Félix and Braendle 2010). This eukaryotic organism provides the opportunity to study genetic mechanisms in the whole animal rather than a cell culture. In addition, it can consume a bacterial or an axenic food source thereby enabling adaptation studies related to diet. Placing the worms from bacteria seeded agar plates (OP50 NGM) to an aquatic axenic media poses a drastic change for the animals. For example, the animals display locomotive differences in the liquid by swimming via a rapid C-shape thrashing motion as opposed to crawling via a sinusoidal configuration on solid (Pierce-Shimomura et al. 2008). Behavior and physiological differences arise when the worms are grown in aquatic axenic media (such as CeMM; *C. elegans* Maintenance Medium or CeHR;*C. elegans* Habitation and Reproduction) CeMM (Szewczyk et al. 2003; Nass and Hamza 2007). In axenic media, the worms exhibit phenotypical alterations such as delayed development, reduced fecundity, increased longevity, and altered body morphology (Houthoofd et al. 2002; Szewczyk et al. 2006; Doh et al. 2016). Environment changes can trigger different gene expression profiles and these profiles can be transmitted to the subsequent generations (Daxinger and Whitelaw 2010; Jaenisch and Bird 2003; Heard and Martienssen 2014). Changing the environment of the animals from agar to axenic media, therefore, can cause transcriptome-level alterations which may have a sustained effect on the progenies.

In its natural habitat, *C. elegans* occupies microbiota-rich decaying plant material and experiences a continuous environment shift between aquatic and terrestrial conditions along with the variations in its diet (Félix and Braendle 2010). The commonly used wild-type laboratory strain of *C. elegans* (N2) has been cultured mainly on bacteria-seeded agar plates for over 40 years (Brenner 1974). It is largely unknown if the domestication of the N2 strain maintained in the laboratory environment has created a selective pressure on the animals (Sterken et al. 2015). Some of the essential questions that await answers are whether: 1.) the adaptive responses depend on the genotype or the environment change; 2.) the environment change-induced gene expression profiles are transmitted to the successive generation in equal levels for different genotypes; 3.) the environment change-induced gene expression profiles are transmitted to the next generation equally for different environmental conditions.

To study the transcriptomic responses in the adaptation of *C. elegans* from terrestrial to aquatic environments, we maintained the P_0_ worms on bacteria-seeded agar plates and F_1_ in liquid cultures. By culturing F_2_ generation back on agar, we identified the maintained transgenerational transcriptional responses. The domesticated N2 strain and a wild isolate AB1 strain were used to examine if a different genotype attributes to the transcriptional dynamics. We revealed substantial variances in gene expression profiles resulting from different genotypes, environments, and genotype-environment interactions. Our results from RNA-sequencing (RNA-seq) data analysis and microscopy showed consistency for the predicted and observed phenotype. Furthermore, we demonstrated that a large number of genes are expressed only in specific growth conditions and many of these genes previously lacked experimental expressed sequence tag (EST) or cDNA evidence.

## Results

To induce an adaptive response and observe its sustained effects on the following generation, the P_0_ generation *C. elegans* was grown on agar, the F_1_ generation in liquid, and the F_2_ generation back on agar (Supplemental Fig. S1). We carried out three sets of experiments with different food sources and animal strains. For the first experiment, the F_1_ generation N2 strain was grown in axenic CeHR Medium. To examine the effect of dietary changes in the liquid cultures, we cultured F_1_ generation N2 strain in bacterial S-Medium for the second experiment. Finally, the adaptive response differences were investigated by growing F_1_ generation AB1 strain in CeHR Medium. For each generation, RNA-seq was performed on young adult animals with three replicates to determine the transcriptional responses (Supplemental Table S1).

### Distinct Gene Groups are Expressed Exclusively in Specific Environmental Conditions

The transcriptome-based clustering of the experiments in the dendrogram of Jensen-Shannon distances demonstrated that the genotypes are separated from each other (Fig. 1*A*). Exposure to different environmental conditions causes the domesticated N2 strain genes to be separated even further than the AB1 genes as illustrated by the dendrogram (Fig. 1*A*) and principal component analysis (Fig. 1*B*). These results suggest that the genotype plays the main role in adaptive response, and the domestication of the animals can cause highly distinct transcriptome profiles under environment changes. We categorized the genes by using Venn diagrams to identify if their expressions are condition specific (Fig. 1*C*). We considered the genes expressed if their Fragments Per Kilobase of transcript per Million mapped reads (FPKM) values are greater than one. The majority of the expressed genes are conserved in the P_0_, F_1_, and F_2_ generations. Nevertheless, hundreds of genes are expressed only under particular conditions. The numb of the uniquely genes expressed in CeHR or S-Medium varies among the experiments (Chi-squared test with Yates correction; P-value < 0.01). Interestingly, the number of genes kept in the epigenetic memory after exposure to an environment change (F_1_F_2_) showed significant difference between the AB1 and N2 strains, but not within the N2 strain experiments (Chi-squared test with Yates correction; P-value < 0.01 and P-value = 0.2, respectively). Together, these results suggest that the condition-specific expression of genes are independent of the genotype and the environment separately—but dependent on their interaction, whereas the formation of the epigenetic memory depends on the genotype.

**Figure 1.**
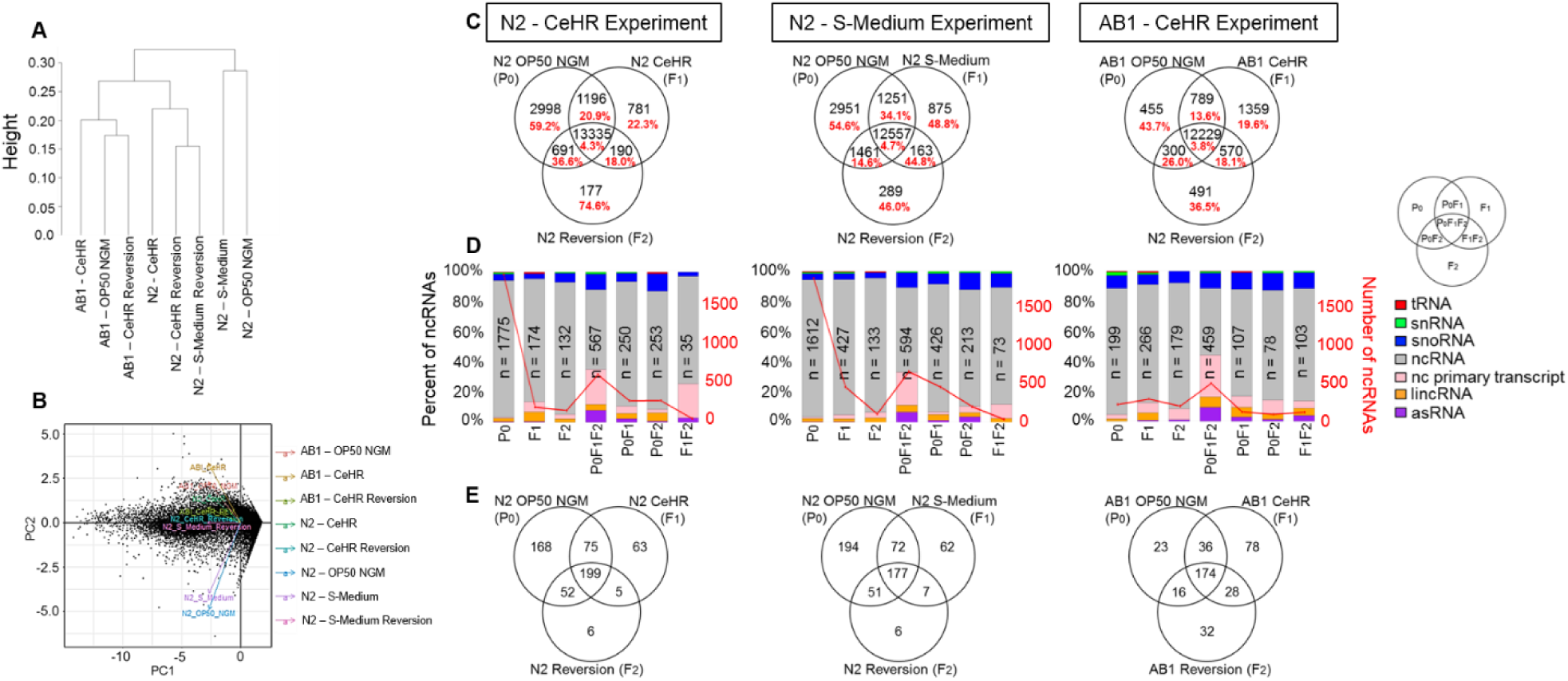
Different strains and environmental conditions present variations in transcriptional responses. **(A)** Dendrogram created based on the gene expression (FPKM) values for the strains and the environmental conditions. **(B)** Principle component analysis plot of gene expression data. **(C)** Categorization of the expressed genes in the three experiments. Red labels represent the percentages of ncRNAs. **(D)** The barplots demonstrate ncRNA profiles in the three experiments. **(E)** Categorization of the previously unconfirmed genes that expressed in our experiments.

Further analyses revealed that non-coding RNA (ncRNA) molecules are enriched (20% -75%) in our pool of condition-specific transcripts indicating their function in genetic adaptation (Fig. 1C). To note, only the polyadenylated ncRNA molecules were included in the analysis and piRNA and miRNA molecules were excluded from the analysis due to their short lengths. The ncRNA molecules were predominantly observed more in the P_0_ generation of N2 strain compared to the P_0_ AB1 strain. Moreover, lower number of ncRNA molecules were expressed when the animals are exposed to an environment change (Fig. 1C, D, and Supplemental Fig. S2). Our results indicate that ncRNA expression is mainly silenced when the worms are exposed to an environment change. The fact that a laboratory condition presents an environment change for the wild-isolate can explain the reason why P_0_ animals of AB1 showed less ncRNA expression compared to the N2 strain. One particular exception to this pattern is the ncRNA molecules common to all of the experiments. We identified that the majority of these ncRNAs (54% - 70%) are conserved among the three experiments (Supplemental Dataset S2). This conservation implies that this group of ncRNAs plays important regulatory functions in the cell and their expression is required regardless of the environment changes.

We reasoned that if a particular group of genes is only expressed under specific conditions, we might detect transcript evidence to previously experimentally unconfirmed genes, especially considering that CeHR is not a common media used in *C. elegans* research. As expected, many unconfirmed genes showed expression in response to the CeHR environment. Particularly, we were able to identify transcript evidence to 21% of the list of unconfirmed genes from WormBase (Harris et al. 2014) by simply growing the animals in different environmental conditions. Similarly with the confirmed genes, the epigenetic memory formed after F_1_ generation demonstrated a difference between the strains (Chi-squared test with Yates correction; P-value < 0.001), but not within the N2 strain.

### Environment, Genotype, and Environment-Genotype Specific Expression of Genes

We grouped the genes based on their genotype, environment, or genotype-environment interaction specific expression (Fig. 2 and Supplemental Dataset S1). For this analysis, a more stringent criterion was used by including the genes with expression levels higher than *pmp-3* of FPKM = 21. This housekeeping gene was selected as a reference because of its stable expression levels found both in previous studies and our analysis (Hoogewijs et al. 2008; Zhang et al. 2012). We defined the genes detected in only one environmental condition for both the strains as “environment-specific”, and found that CeHR-specific gene expression displayed Gene Ontology (GO) enrichment for the neuropeptide signaling pathway. Given that this pathway functions in numerous behavioral activities such as reproduction, locomotion, mechanosensation and chemosensation, it is likely that neuropeptide signaling pathway plays a key role in adaptation to CeHR and is evolutionarily conserved between the strains (Li and Kim 2008).

**Figure 2.**
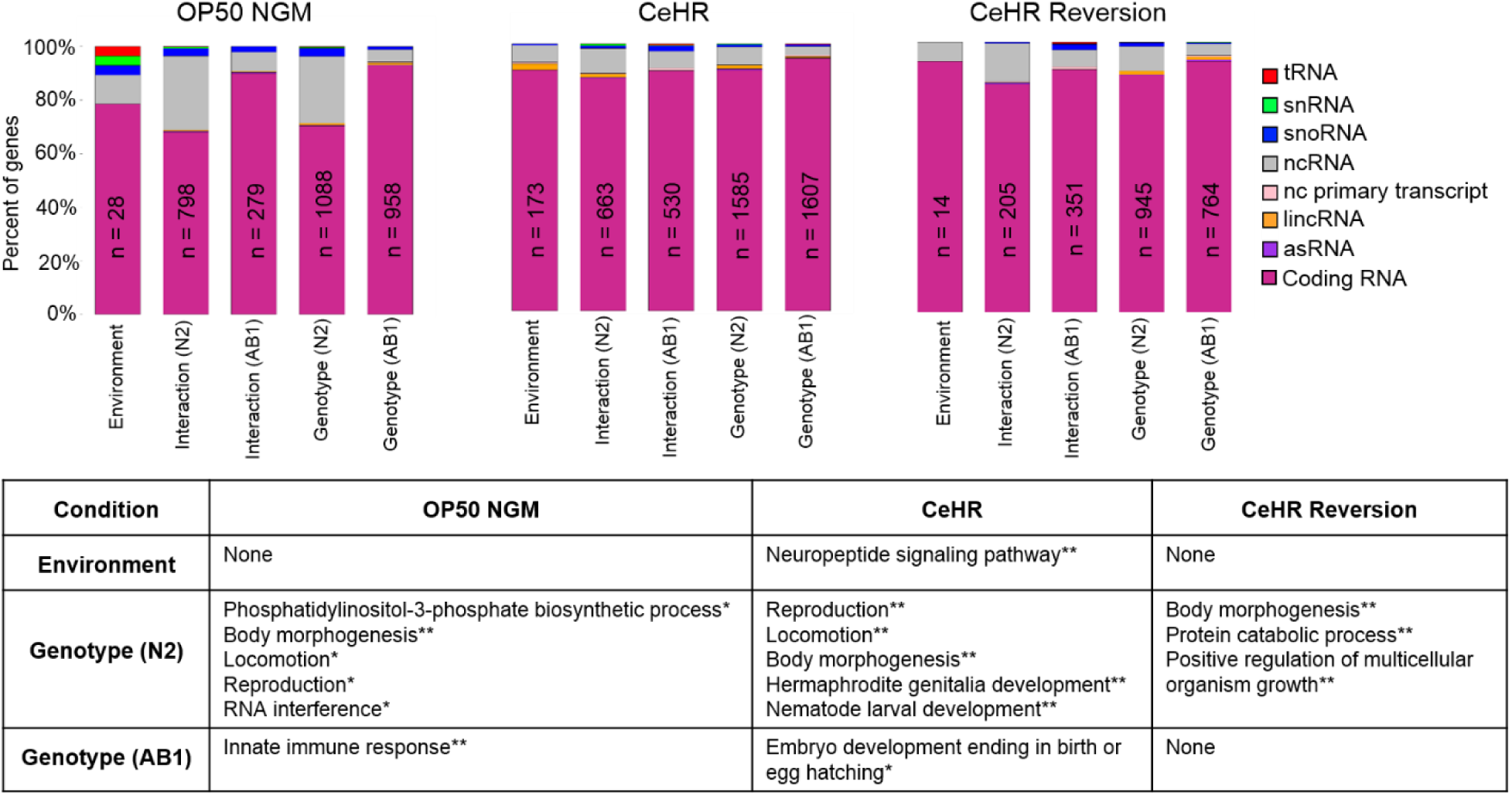
Shown are the number of gene expressions exclusive to environment, genotype, or genotype-environment interactions. The number of the expressed ncRNAs are decreased under stress conditions. The table depicts the enriched gene ontology (GO) terms assigned to environment or genotype specifically expressed genes. Genotype-environment interaction genes did not demonstrate enrichment for GO terms. “None” represents the groups with no GO enrichment. Benjamini Hochberg corrected P-values for the GO terms: ^*^ < 0.01; ^**^ < 0.001.

The “genotype-specific” expression represents the genes found in only one of the strains under a single environmental condition. All the N2 specific genes showed GO enrichment for body morphogenesis in all the three generations while reproduction and locomotion GO enrichment were observed in P_0_ and F_1_ generations (Fig. 2). These genes may be the ones that have undergone genetic assimilation in response to the domestication of the animals. Previously we reported that the N2 worms produce less progeny compared to the AB1 animals in CeHR (Doh et al. 2016). Our analysis showed that along with the reproduction genes, genes related to hermaphrodite genitalia development were expressed exclusively in N2 strain in CeHR. This finding may suggest vulva aberrations in the reproductive system. In addition, the exclusive expression of the genes functioning in positive regulation of multicellular organism growth may help the animals readapt to the OP50 NGM condition in the F_2_ generation of the N2 animals.

The genes expressed solely under a particular environment and strain were categorized as “genotype-environment interaction” genes (Fig. 2). This group of genes did not demonstrate an enrichment of GO terms. The proportion of ncRNA molecules in our group of genes showed correlation with our hypothesis that the majority of the ncRNAs are silenced under changes in environment conditions. For example, the N2 worms showed relatively higher proportions of ncRNAs for interaction and genotype-related genes on OP50 NGM, but these proportions were much lower in more stressful CeHR and CeHR reversion (Fig. 2).

### Domesticated Strain Presents Higher Differential Expression under Environment Changes

To investigate the differences in transcriptional responses between the strains and among the environmental conditions, we identified the differentially expressed genes (DEGs). The estimated fold change responses of ten randomly selected transcripts were confirmed by quantitative RT-PCR (Supplemental Fig. S3 and Supplemental Table S2). We found the highest number of DEGs between OP50 NGM and CeHR conditions for the N2 strain indicating that the change in diet and physical environment triggers more transcriptional responses for adaptation of the domesticated strain (Fig. 3). The number of DEGs were much lower for the F_2_ generation of the same experiment, indicating that the majority of the gene expressions started to resemble the P_0_ animals. As most of the gene expression patterns return to the steady-state levels after a short period of time, this finding was expected (López-Maury et al. 2008). The change in only the physical condition introduced by the S-Medium apparently did not cause as drastic transcriptional responses as the CeHR since the number of the DEGs were lower. However, S-Medium reversion conditions demonstrated higher DEGs compared to OP50 NGM. We note that we are able to maintain the worms in S-Medium for only one generation, potentially because of the toxins released by the bacteria in the aquatic environment (Gems and Riddle 2000; Croll et al. 1977). Therefore, additional genetic mechanisms may be required for re-adaptation of the animals from S-Medium. Transcriptional variations were more modest across the generations for AB1 animals.

**Figure 3.**
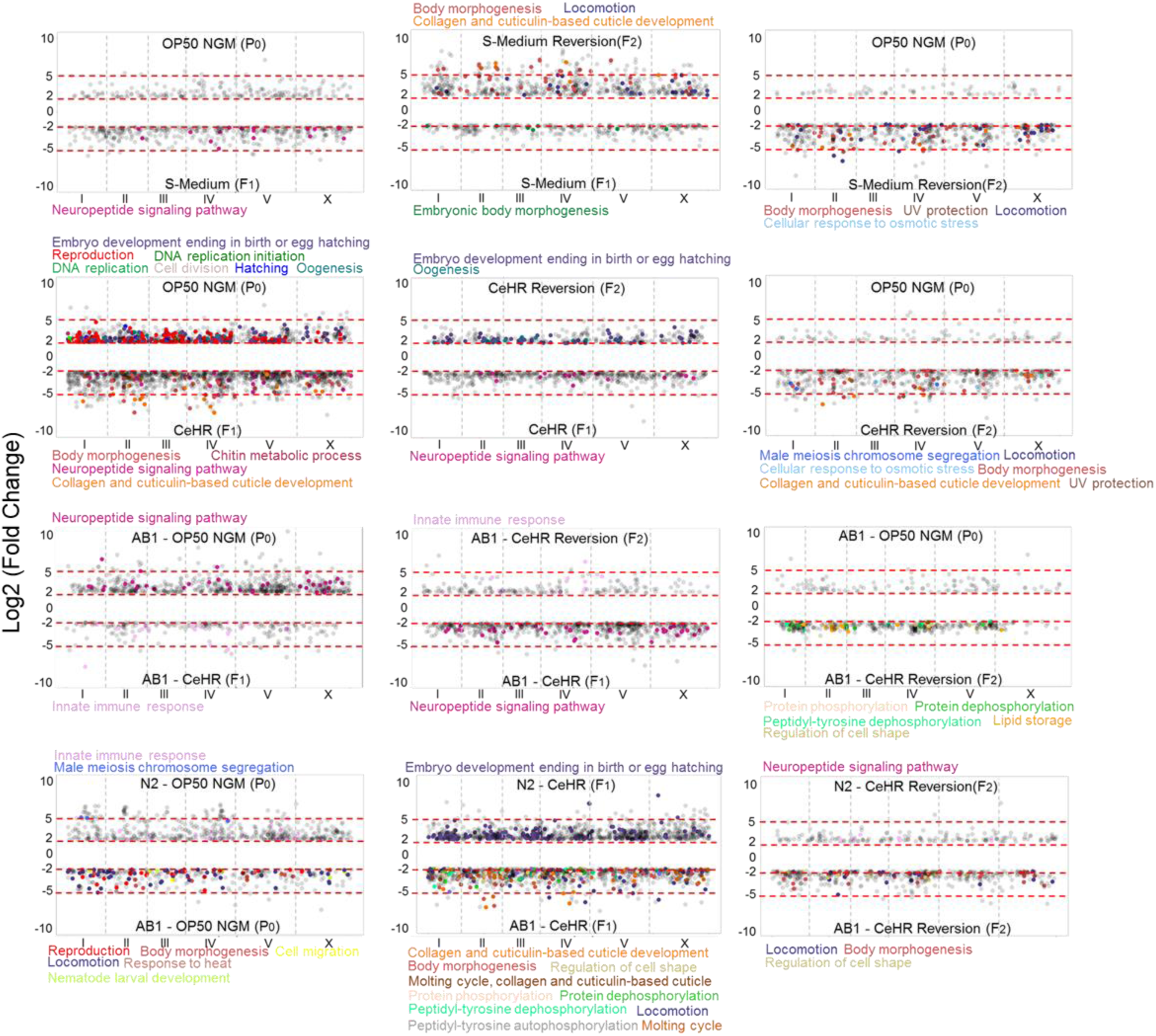
The differential expression of the genes between the environment conditions and the strains. The enriched GO terms for the genes were represented with the colors. P-value < 0.05 for all the enriched GO terms. X-axis represents the chromosomal regions.

Particular environmental change induced DEGs in F_1_ maintained their upregulation or downregulation in the next generation. The transmission of the DEGs to the F_2_ generation from F_1_ was between 20% - 34% (Fig. 3). We hypothesized that if these genes are indeed passed to the next generation through epigenetic memory, they should present sequence motif enrichment in their promoter region to enable their recognition. Intriguingly, the genes kept in the epigenetic memory after exposure to CeHR showed enrichment for different sequence motifs (Fig. 4*A*). This difference was expected as the DEG profiles were highly variant between the strains (Fig. 3). Similarly, the DEGs in the F_1_ generations presented enrichment on distinct tissues (Fig. 4*B*). Altogether, these findings suggest that the domestication of the animals causes significant alterations in the adaptive responses, and the environmental changes affect the transcriptomic response in a unique manner.

**Figure 4.**
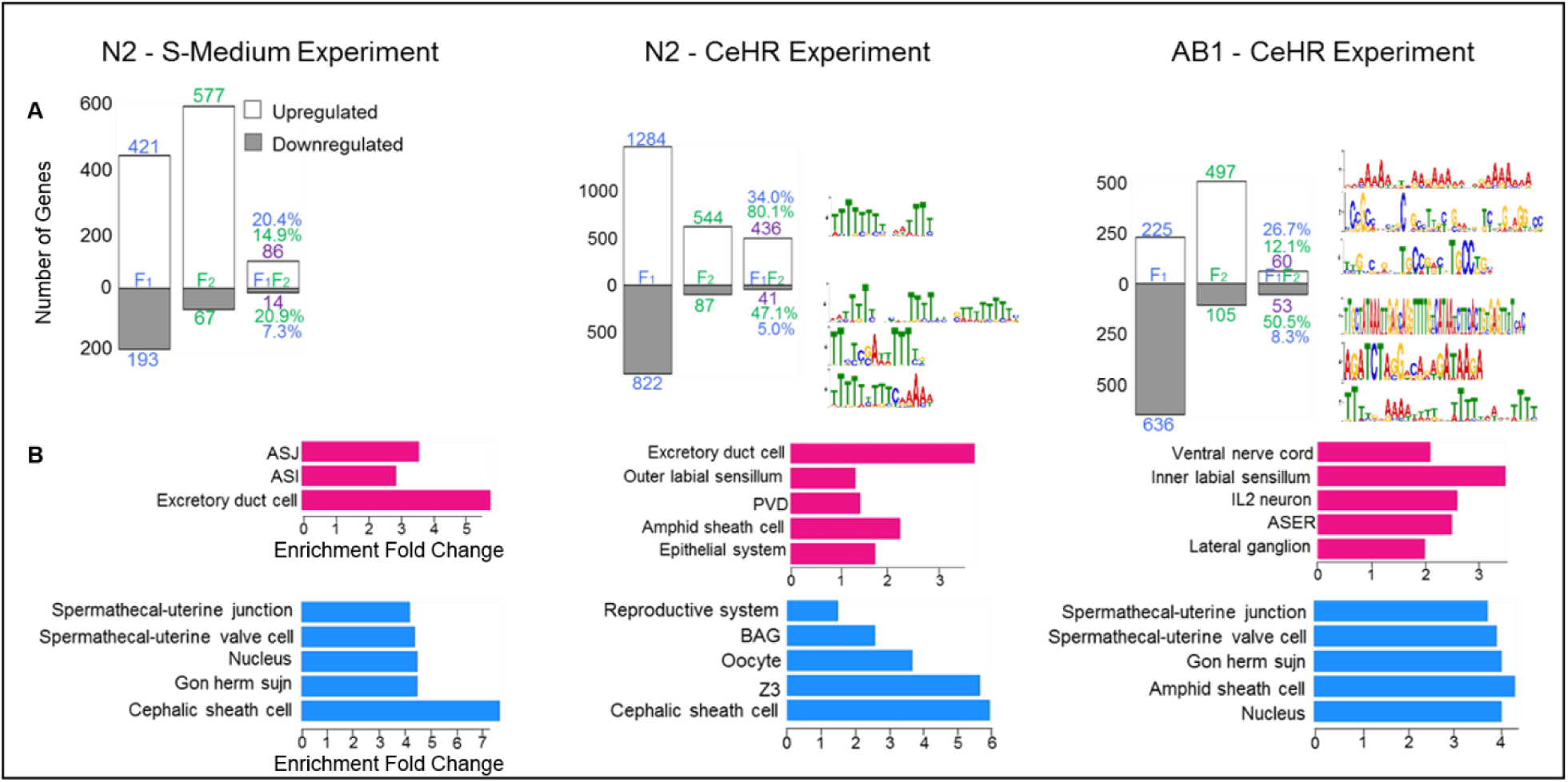
Different strains and the environment conditions contribute to differences in the multigenerational gene expression profiles. **(A)** Percentage of the DEGs in the F1 generations transmitted to the F2 generations (see the third category – F1F2). The DEGs in the F1 and F2 generations were identified in comparison to the gene expressions in the P0 animals. The gene expressions transmitted through the epigenetic memory after exposure to CeHR showed distinct enrichments for sequence motifs in their promoter regions for N2 and AB1 strains. **(B)** Tissue enrichment of the DEGs under exposure to environmental changes in F1 generation.

CeHR-grown N2 and AB1 *C. elegans* have been shown to display different phenotypical traits on their body morphology (Doh et al. 2016). In accordance with this finding, the GO enrichment analysis results demonstrated enrichment for body morphogenesis for upregulated genes in CeHR compared to OP50 NGM in the N2 strain (Fig. 3). The fact that this GO term is not enriched for AB1 implies that the phenotype is caused by a separate transcriptional mechanism.

Another interesting finding is that neuropeptide signaling pathway genes were enriched in response to environment changes. The pathway genes were upregulated in the aquatic environments compared to agar in N2 and in agar compared to the aquatic environment in AB1. Considering that the natural habitat of *C. elegans* is decaying material rather than soil, placing the wild-isolate animals onto agar plates can create a more stressful environment to the animals than CeHR (Félix and Braendle 2010; Sterken et al. 2015). These findings support our hypothesis that neuropeptide signaling pathway is a key mechanism in adaptation and it is conserved in different strains of *C. elegans*.

### Differentially Expressed Gene Profiles Show Correlation with the Observed Phenotype

We wondered if the functional data analysis results indeed points to phenotypical alterations in the animals. To test this idea, we employed microscopy techniques to detect the potential phenotypes. In the N2 strain, the genes functioning in reproduction and oogenesis have been downregulated in CeHR compared to agar and reversion. In AB1, the DEGs among the conditions did not show the same GO enrichment. This pattern is an indicator of phenotypical changes in the reproductive system that can be observed in CeHR but absent in reversion for N2. To test our prediction, the germlines of N2 and AB1 adults in the different growth conditions were examined via confocal microscopy (Fig. 5). CeHR grown N2 but not AB1 adults displayed aberrant germlines, suggesting that the reduced fecundity CeHR raised N2 animals may have resulted from the inability of animals to make gametes. In agreement with our hypothesis, the worms placed back to agar plates did not exhibit the similar phenotypical abnormalities with CeHR grown animals (Supplemental Fig. S4).

**Figure 5.**
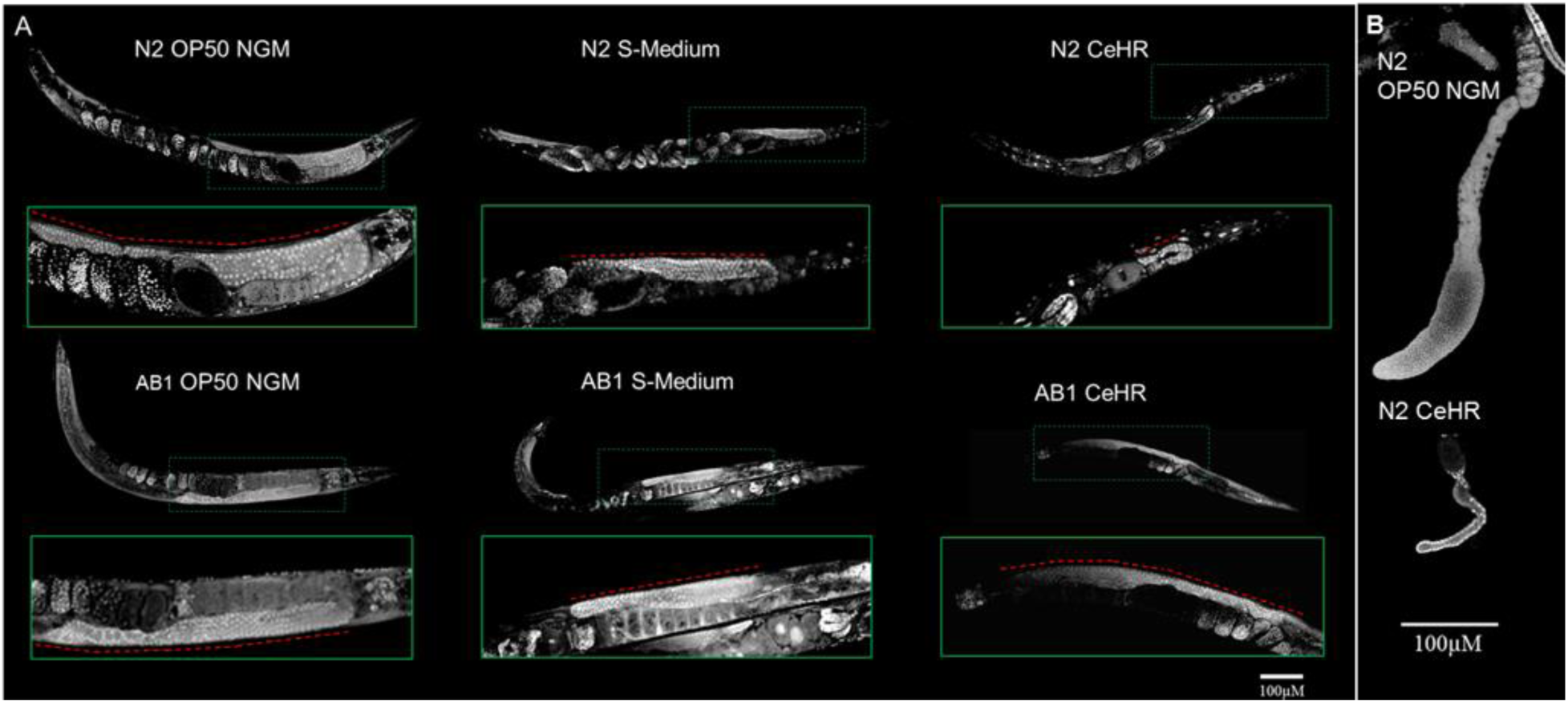
N2 but not AB1 *C. elegans* exhibit germline abnormalities in CeHR. **(A)** N2 and AB1 germlines exhibit normal germ cell morphologies and numbers in OP50 NGM. Germlines of S-Medium grown animals are similar to OP50 NGM grown animals. However, embryos accumulate in S-Medium grown adults. As expected, N2 adults grown in CeHR exhibit aberrant germlines with drastically reduced germ cell numbers, which results in an overall smaller germline. N2 adults in CeHR have significantly fewer number of oocytes and a high number of internally hatched larvae. These abnormalities are not observed in CeHR grown AB1 adults. **(B)** Dissected out gonadal arms of N2 *C. elegans* grown on OP50 NGM and CeHR.

Then, we asked if these germline abnormalities were caused by an increase in physiological germ cell apoptosis. We utilized a transgenic line of *C. elegans* that expresses functional CED-1::GFP fusion protein in sheath cells (Zhou et al. 2001). CED-1 functions by initiating a signaling pathway in phagocytic cells that promotes cell corpse engulfment and is commonly used as a marker to detect cells undergoing the apoptotic pathway. In CeHR grown adults, CED-1::GFP was detected in a much higher proportion of germ cells (Fig. 6*A*). Why the number of apoptotic germ cells in CeHR grown N2 adults were increased and whether or not this was the sole cause of aberrant germlines will require further investigations.

**Figure 6.**
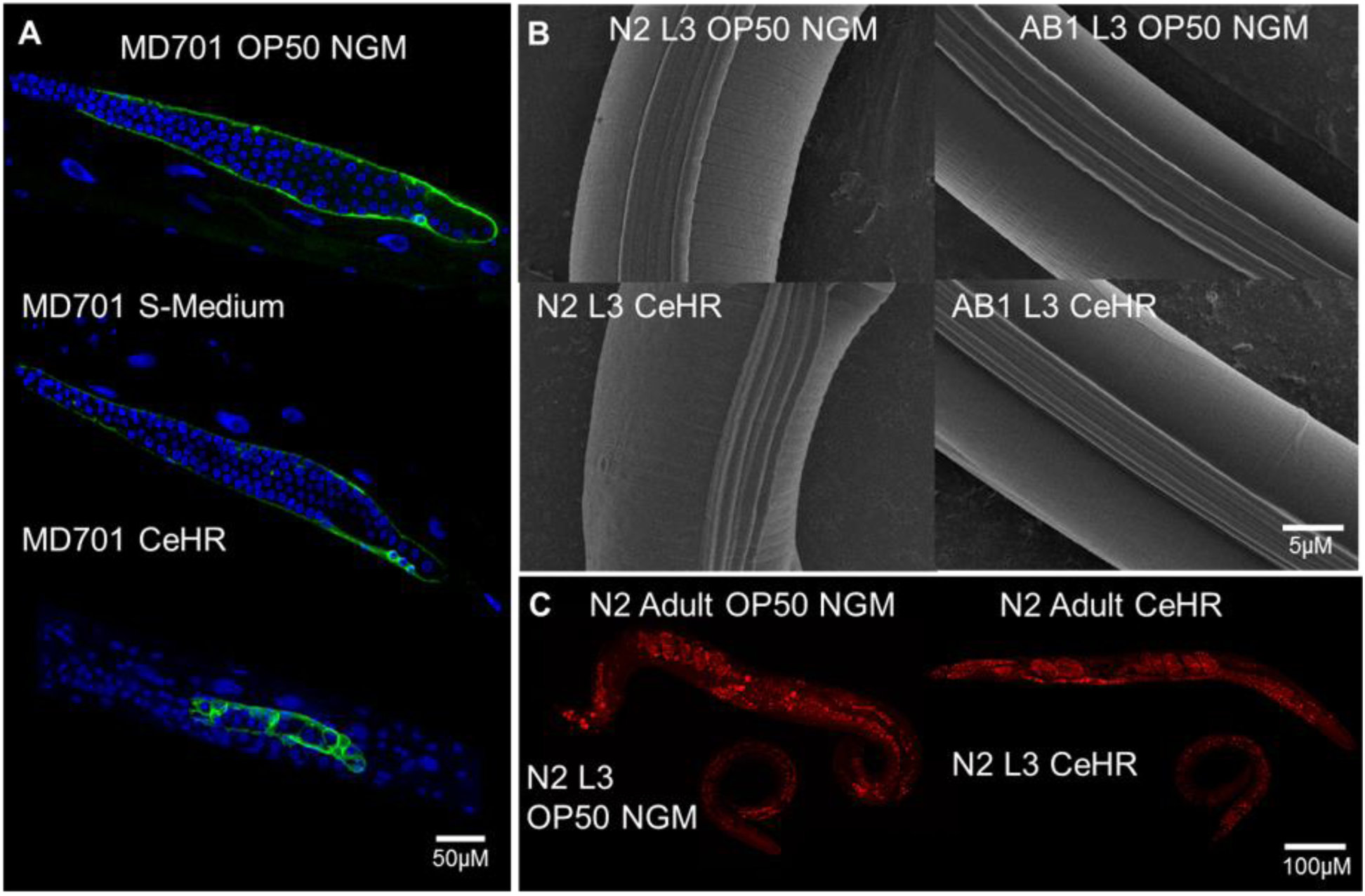
*C. elegans* grown in CeHR exhibit higher number of physiological germ cell apoptosis, more pronounced alae structures, and smaller fat stores. **(a)** OP50 NGM, S-Medium and CeHR grown MD701 adults were stain with DAPI. One or two germ cells enclosed by CED-1::GFP are detected around the loop region of the gonads in MD701 adults grown in OP50 NGM and S-Medium. CED-1::GFP is detected around a large proportion of germ cells in CeHR grown MD701 adults. **(b)** Scanning electron microscope (SEM) micrographs of OP50 NGM and CeHR grown N2 and AB1 L3 animals. N2 and AB1 L3 animals display normal alae and annuli with grown on OP50 NGM. N2 but not AB1 L3 exhibit more pronounced alae in CeHR. **(c)** Nile red staining of L3 and adult N2 *C. elegans* grown on OP50 NGM and CeHR. The number of fat stores are indistinguishable between OP50 NGM and CeHR grown N2 animals, but the fat stores significantly larger in OP50 NGM than CeHR raised N2 adults around the abdominal and tail regions.

For the N2 worms, another enriched GO term was collagen and cuticulin-based cuticle development for upregulated genes in CeHR compared to agar and reversion conditions. Hence, we reasoned that the cuticles should show morphological differences between agar and CeHR-grown N2 animals. In adult animals, cuticular structures were difficult to distinguish between N2 and AB1 via scanning electron microscopy (SEM). More pronounced cuticular structures were detected in the earlier larvae stages with SEM. We observed more protruding alae structures in CeHR grown N2 larvae compared to AB1 larvae, but the annuli were indistinguishable between the two strains (Fig. 6*B*). Taken together, the correlation between the function of the DEGs and the observed phenotype suggests that these genes are potentially a part of underlying genetic networks causing these phenotypes.

In our previous study, we demonstrated that the body morphology of CeHR grown animals was thinner and longer than OP50 NGM raised animals (Doh et al. 2016). We asked if this change in body morphology was due to a reduction in fat stores that may have resulted from the absence of a bacterial diet. N2 animals grown on OP50 NGM and CeHR were stained with Nile Red (Greenspan et al. 1985) to visualize fat stores in all tissues of L3 and adult animals. We observed no significant difference in the number of fat stores between OP50 NGM and CeHR-grown N2 animals; however, larger fat stores were apparent in the abdominal and tail regions of OP50 NGM grown N2 adults (Figure 6*C*). Thus, the lack of a bacterial diet apparently contributes to the morphological differences through the fat storage.

## Discussion

In this paper, we demonstrated that drastic environment changes cause significant alterations in the transcriptional responses in a *C. elegans* model. Domestication of the worms leads to unique transcriptional characteristics in the adaptive responses, and these characteristics show correlation with the observed phenotypes. For instance, we detected differential expression in the body morphology-related genes along with the phenotypical changes on the body morphology in CeHR-grown N2 strain, but not the AB1 strain. Furthermore, we found that different environments provoke the condition-specific expression of genes, and some of these genes lacked previous experimental EST or cDNA evidence.

Along with the phenotypical differences, there were considerable differences in RNA expressions between the N2 and AB1 strains which were grown in the same growth conditions. Previous studies have determined that the N2 and wild-isolate CB4856 strains display a high variation in gene expressions and many of these strain-specific variations are related to innate immunity genes (Capra et al. 2008). Similarly, our GO analysis results showed enrichment in innate immune response for the genes expressed exclusively in the AB1 strain on OP50 NGM (Fig. 2). The AB1 animals are challenged more with pathogens in their natural habitat. The AB1-specific expression of innate immune response genes indicates that the expression of these genes was kept in the epigenetic memory to prepare the animals for a potential pathogen encounter. This expression profile is lost in domesticated N2. Collectively, our results suggest that *C. elegans* in the wild have the natural aptitude to survive not only on different food sources but also many other environmental variables including pathogens or the shifts between the terrestrial and aquatic settings. However, years of laboratory domestication of the N2 strain may have given rise to laboratory selections or genetic bottlenecks that may have hindered the ability of the N2 animals to acclimate to changing environments (Sterken et al. 2015).

Our results revealed that specific groups of genes are expressed only in particular environments. Some of these genes lacked transcript evidence previously (Fig. 1*D*). Considerable effort has been devoted to annotating the genes in *C. elegans*, but the standard WormBase (Harris et al. 2014) annotation still contains thousands of “predicted genes” (about 8%) which lack direct experimental cDNA or EST evidence (Gerstein et al. 2010). The predicted genes in the *C. elegans* genome have been largely identified by computational methods. Experimental detection of these genes was missing mainly because they demonstrate poor expression, weak statistical signals or less conservation across species (Hillier et al. 2005). Others found that unused genetic information can remain in the genome for many generations (Collin and Cipriani 2003). Reactivation of dormant genes has been used for the treatment of particular disorders indicating that the silenced genes can have important functions in the genome (Huang et al. 2011). These studies provide an evidence that dormant genes can be available on the genome throughout many generations and they can contain essential genetic information. Thus, we entertain the possibility that these predicted genes simply need a trigger to activate their expression, and this trigger may be elicited when animals are exposed to certain environment stimuli.

A better understanding of the biological systems can be achieved through mining publicly available data sources (Çelen et al. 2015). We expanded our knowledge in the affected genetic systems by acquiring the ncRNA profiles from WormBase (Harris et al. 2014). Condition-specific expression of ncRNAs implies important regulatory functions for these molecules in adaptation. Earlier studies have identified various classes of ncRNAs and their functions in biological systems including RNA splicing, DNA replication, epigenetic regulation of gene expression, and X-chromosome inactivation (Carthew and Sontheimer 2009; Kaikkonen et al. 2011; Kishore 2006; Zhang et al. 2011; Cao 2014). However, the majority of predicted ncRNAs have properties or functions that have not been identified yet. The list of unclassified ncRNAs, or “ncRNAs,” employed in this study was derived from the ‘7k-set’ that was generated by the modENCODE Consortium (Gerstein et al. 2010). The ‘7k-set’ was assembled via predictions based on conservation and RNA secondary structure, and therefore, functional genomic studies of these “ncRNAs” are also still lacking. The co-enrichment of the ncRNA expressions with the coding ones suggest an interplay between these two.

In our RNA-seq experiments, we used poly(A) selection method but still observed numerous ncRNA molecules. In fact, many eukaryotic ncRNAs are polyadenylated. For example, a poly(A) tail is part of the mature RNA for many long ncRNAs, i.e. *Xist* that mediates X-chromosome inactivation (Amaral and Mattick 2008). We cannot rule out that the ncRNAs sampled in our study resulted from technical artifacts in the RNA-seq. Nevertheless, many ncRNAs had expression levels higher than the housekeeping gene *pmp-3*. These findings would seem to suggest that the sampling of these RNAs was not an artifact of RNA-seq. The inclusion of non-polyadenylated ncRNA molecules can bring more insights for understanding the ncRNA functions in adaptation.

For the majority of the *C. elegans* studies, the worms are cultured on bacteria-seeded agar plates instead of an axenic medium (Stiernagle 2006; Sulston and Hodgkin 1988). However, a bacterial food source can present confounding effects on certain studies. For example, bacterial by-products can create genetic responses in the animals and these responses are often overlooked. Axenic media is desirable in space, biochemical, and toxicology case studies as it enables automated culturing and experimentation, and it eliminates the potential contamination risks due to a bacterial diet (Adenle et al. 2009; Doh et al. 2016; Samuel et al. 2014). Placing the animals in axenic media, however, have profound consequences altering a variety of biological processes. To distinguish whether the genetic responses are from the case study or the axenic media conditions, a separate investigation should be conducted. In light of this, we revealed that *C. elegans* acquire large variations in gene expressions upon single generation exposure to two types of liquid media. We believe our data can provide standard controls for future studies that utilize liquid cultivation of *C. elegans* for experimentations.

## Methods

### C. elegans strains and growth conditions

Wild-type N2, wild-isolate AB1, and CED-1::GFP (MD701) strains were obtained from the Caenorhabditis Genetics Center (CGC). All animals were grown at 21^°^C. Embryos were isolated via the bleaching protocol (Sulston and Hodgkin 1988) and placed in the following growth media: *E. coli* OP50 seeded on NGM agar plates (OP50 NGM), 1 mL of S-Medium in 20 mL scintillation vials inoculated with a concentrated *E. coli* OP50 pellet made in 6 mL of an overnight culture (OP50 S-Medium), and CeHR medium in 20 mL scintillation vials according to Nass and Hamza (Nass and Hamza 2007) with minor modifications.

### RNA Isolation, Illumina Sequencing

Total RNA from young adult *C. elegans* (4 hours post L4) were isolated with a modified TRIzol protocol and recovered by alcohol precipitation. Total RNA was further purified by PureLink (tm) RNA Mini Kit (Life Technologies). Two micrograms of total RNA were used for library preparations using the TruSeq Stranded mRNA LT Sample Prep Kit (Illumina). Libraries were sequenced on an Illumina HiSeq 2500 instrument set to the rapid run mode using single-end, 1 × 51 cycle sequencing reads as per manufacturer’s instructions.

### qRT-PCR

Quantitative RT-PCR was conducted using 1st strand cDNA synthesized from total RNA and gene-specific primers (Supplemental Table S3). Each cDNA sample was amplified using the SYBR® Premix Ex Taq ^(tm)^ II (Takara Bio) on the ABI 7500 Fast Real-time PCR System (Applied Biosystem) according to the manufacturers. The experiment was performed by three independent experiments with biological triplicates.

### Bioinformatics Analyses

The Tuxedo pipeline (Trapnell et al. 2012) was used to find DEGs with default parameters as described in the cited article (see the companion paper). Briefly, quality of the sequencing reads were evaluated with FASTQC (version 0.11.2). The reads were mapped to the WBCel235 reference genome from Ensembl (Cunningham et al. 2014) by using TopHat (version 2.1.1) (Kim et al. 2013). Differential expression analysis was performed with Cuffdiff (version 2.2.1).We considered genes with FPKM > 1, FDR adjusted *p-values* < *0.05*, and log^2^ fold change > 2 as differentially expressed. miRNA, piRNA, and rRNA molecules were discarded from the analysis. The ncRNA molecules (WS250) and unconfirmed genes were acquired from WormBase through the ftp site and personal communications, respectively (Harris et al. 2014). The motif enrichment analysis was performed with MEME Suite (v.4.11.1) (Bailey et al. 2009). The motifs were ranked based on the E-value significance. The promoter sequences for the motif enrichment were retrieved from UCSC Genome Browser database for ce11 (Rosenbloom et al. 2015). GO and tissue enrichment analyses were made by using the Database for Annotation, Visualization and Integrated Discovery (v6.8) and WormBase, respectively (Huang et al. 2008; Angeles-Albores et al. 2016). Dendrogram and principle component analysis plots were generated using R CummeRbund package (Goff et al. 2012). Chi-squared test with Yates correction was applied to find if the number of DEGs show difference between the two strains or between the environment conditions. P-value < 0.05 was considered statistically significant. R software was used for all statistics analyses and the plots (R Development Core Team 2017).

### Microscopy

Nematodes were fixed in cold (-20^°^C) methanol for 15 minutes and stained with SYBR Gold (1:500,000 dilution in PBS) or DAPI (100 ng/mL) for 15 minutes, or Nile Red (5 ng/mL) for 30 minutes, at 21^°^C with gentle rocking. Worms were destained in PBST, washed three times with M9 and mounted on a 2% agarose pad for microscopy. All micrographs were taken with the Zeiss 710 laser scanning confocal microscope with a 40×/1.2 N.A. water immersion objective. Dissected gonads were stained with SYBR gold as mentioned above. To visualize cuticular structures via cryo-SEM, nematodes were placed onto 0.22 μM filters where the excess liquid was removed. Upon attaching the filter paper onto the sample holder, samples were plunged into liquid nitrogen slush. There a vacuum was pulled allowing sample transfer to the Gatan Alto 2500 cryo chamber at a temperature of -125^°^C. Samples were sublimated for 10 minutes at 90^°^C followed by cooling to -125^°^C. A thin layer of Gold Palladium was sputtered onto the samples. The samples were then transferred into a Hitachi S-4700 Field-Emission Scanning Electron Microscope for imaging.

## Data Access

All the sequencing data generated for this study have been submitted to the NCBI Gene Expression Omnibus (GEO; http://www.ncbi.nlm.nih.gov/geo/) under accession number GSE103777.

## Acknowledgements

We thank Dr. Sevinç Ercan for critically reviewing the manuscript and providing helpful comments. This work was supported by NASA grants NNX12AR59G, NNX10AN63H, and NNX13AM08G. The BIOMIX computing cluster and the Bioimaging centers are supported by Delaware INBRE grant (NIH/NIGMS GM103446) and Delaware EPSCoR grants (NSF EPS-0814251 and NSF IIA-1330446). İ.Ç. acknowledges the University of Delaware for Graduate Fellow Award and Summer Doctoral Fellowship.

